# Prediction of the receptorome for the human-infecting virome

**DOI:** 10.1101/2020.02.27.967885

**Authors:** Zheng Zhang, Sifan Ye, Aiping Wu, Taijiao Jiang, Yousong Peng

**Affiliations:** College of Biology, Hunan Provincial Key Laboratory of Medical Virology, Hunan University, Changsha 410082, China; Center for Systems Medicine, Institute of Basic Medical Sciences, Chinese Academy of Medical Sciences & Peking Union Medical College, Beijing 100005, Suzhou Institute of Systems Medicine, Suzhou, Jiangsu 215123, China

## Abstract

The virus receptors are key for the viral infection of host cells. Identification of the virus receptors is still challenging at present. Our previous study has shown that human virus receptor proteins have some unique features including high N-glycosylation level, high number of interaction partners and high expression level. Here, a random-forest model was built to identify human virus receptorome from human cell membrane proteins with an accepted accuracy based on the combination of the unique features of human virus receptors and protein sequences. A total of 1380 human cell membrane proteins were predicted to constitute the receptorome of the human-infecting virome. In addition, the combination of the random-forest model with protein-protein interactions between human and viruses predicted in previous studies enabled further prediction of the receptors for 693 human-infecting viruses, such as the Enterovirus, Norovirus and West Nile virus. As far as we know, this study is the first attempt to predict the receptorome for the human-infecting virome and would greatly facilitate the identification of the receptors for viruses.

## Inroduction

Receptor-binding is the first step for viral infection of host cells. Proteins are considered to be the more ideal receptors for viruses due to their higher binding affinity and specificity than carbohydrate and lipid (Baranowski et al., 2001; Casasnovas, 2013; Dimitrov, 2004; Li, 2015; Wang, 2002). Lots of human virus receptors have been identified. In addition, it is not a random process for viruses to choose proteins as their receptors. Previous studies have shown that the proteins that are abundant in the surface of host cells or have relatively low affinity for their natural ligands are the preferred receptors for viruses (Dimitrov, 2004; Wang, 2002). Moreover, based on a collection of 119 mammalian virus receptors, our recent study has further revealed that human virus receptor proteins have higher level of N-glycosylation, higher number of interaction partners in the human protein-protein interaction (PPI) network and higher expression level in 32 common human tissues compared to other cell membrane proteins (Zhang et al., 2019). The results obtained from these studies could facilitate the identification of human virus receptors.

Identification of virus receptors in host cells is challenging. Currently, several experimental methods have been developed for identifying virus receptors. The first approach is to select candidate membrane proteins that can bind to the virus receptor-binding proteins (RBPs) by affinity purification and mass spectroscopy (Free et al., 2009; Ryu, 2016). The second approach is firstly to identify monoclonal antibodies which can block the virus entry, and then take the membrane proteins to which the monoclonal antibodies bind as candidate receptor proteins (Minor et al., 1984; Ryu, 2016). The third approach is to identify virus receptors by functional cloning selection based on the cDNA expression library (Ryu, 2016). The proteins that can enable viral infection of the non-susceptible cells after transfection are considered as the receptor candidates. However, the identification of virus receptors is still time-consuming and difficult at present. Besides, the limitations on scalability have hampered the large-scale identification of viral receptors. Therefore, it is urgent to develop the computational methods for the identification of the human virus receptors.

Previous studies have developed several computational models for predicting the PPIs between viruses and hosts which can help identify virus receptors (Lasso et al., 2019; Yan et al., 2019). For example, Lasso et al. developed an in silico computational framework (P-HIPSTer) that employed the structural information to predict more than 280,000 PPIs between 1,001 human-infecting viruses and humans, and made a series of new findings about human-virus interactions (Lasso et al., 2019). The predicted PPIs between viral RBPs and human cell membrane proteins can be used to identify virus receptors. Here, a computational model was developed to predict the receptorome of the human-infecting virome based on the features of human virus receptors and protein sequences. Furthermore, the combination of this computational model with the PPIs predicted in Lasso’s work was further used to predict the receptors for 693 human-infecting viruses. The results of this study would greatly facilitate the identification of human virus receptors.

## Materials and Methods

### Human virus receptors, human cell membrane proteins and human membrane proteins

A total of 91 human virus protein receptors were obtained from the viralReceptor database (available at http://www.computationalbiology.cn:5000/viralReceptor) that was developed in our previous study (Zhang et al., 2019). Human cell membrane proteins and human membrane proteins were obtained from the UniProtKB/Swiss-Prot database on February 21, 2020. The human proteins with the words of “cell membrane” and “cell surface” in the field of “Subcellular location” were considered to be human cell membrane proteins. The human proteins with the words of “membrane” and “cell surface” in the field of “Subcellular location” were considered to be human membrane proteins. A total of 3,642 human cell membrane proteins and 7,663 human membrane proteins were obtained.

### Features of the human virus protein receptors and protein sequences

The N-glycosylation sites of the human proteins mentioned above were obtained from the UniprotKB/Swiss-Prot database. For those proteins without the annotation of N-glycosylation sites in the UniprotKB/Swiss-Prot database, their N-glycosylation sites were predicted with NetNGlyc 1.0 (available at http://www.cbs.dtu.dk/services/NetNGlyc/) (Gupta et al., 2004). The N-glycosylation level of these proteins was defined as the number of N-glycosylation sites per 100 amino acids.

To calculate the node degree of the human proteins in the human PPI network, firstly, the human PPIs with the combined scores greater than 400 were extracted from the STRING database (version 10.5) (Szklarczyk et al., 2015) and were used to form the human PPI network. Then, the node degree was calculated with the function of *degree* in the R package igraph (version 1.2.4.2) (Csardi and Nepusz, 2006).

The expression level of the human genes in 32 common human tissues was obtained from the Expression Atlas database (Petryszak et al., 2016) on February 6, 2018. Since there were strong correlations between the gene expression level in different tissues, the principal component analysis (PCA) method was used to reduce the correlations with the function of *PCA* in the package scikit-learn (version 0.21.3) (Pedregosa et al., 2011) in Python (version 3.6.7). Only the first principal component was used to measure the expression level of human genes, which explained 95% of the total variance.

The frequencies of k-mers with one or two amino acids were calculated for each human protein with a Python script.

### Machine learning modeling

To distinguish human virus receptors from other human cell membrane proteins using machine learning models, the human virus receptors were chosen as positive samples. Since the human cell membrane proteins may contain virus receptors unidentified yet, the human membrane proteins were taken as negative samples after excluding the human cell membrane proteins.

Because not all human proteins were observed in the used human PPI network or showed the observed expressions in the available data, only the human proteins that possess all the three protein features, i.e., the N-glycosylation level, node degree and expression in common human tissues, were used in the modeling. Besides, the sequence redundancy in both human virus receptor proteins and human membrane proteins was removed using CD-HIT (version 4.8.1) (Fu et al., 2012) at 70% identity level. Finally, a total of 88 human virus receptors and 1,743 human membrane proteins were used in the machine learning modeling.

The random-forest (RF) model is an ensemble machine learning technique using multiple decision trees and can handle data with high variance and high bias, while the risk of over-fitting can be significantly reduced by averaging multiple trees. Therefore, the RF model was chosen to distinguish human virus receptors from other human cell membrane proteins. Since the number of positive samples (human viral receptors) was much smaller than that of negative samples (human membrane proteins), the function of BalanceRandomForest in the package imbalanced-learn (version 0.5.0) (Chen et al., 2004) in Python was used to deal with the imbalanced positive and negative samples with the parameter of n_estimators set to be 100.

Five times of five-fold cross-validations were conducted to evaluate the predictive performances of the RF model with the function of StratifiedKFold in the package scikit-learn in Python. The predictive performances of the RF model were evaluated by the area under receiver operating characteristics curve (AUC), accuracy, sensitivity and specificity.

### Validation of the RF model by ranking the receptor candidates

The predicted PPIs between human-infecting viruses and human were obtained from the database of P-HIPSTer (available at http://phipster.org/) (Lasso et al., 2019) on November 1, 2019. A total of 9,395 pairs of interactions between viral RBPs and human cell membrane proteins with the likelihood ratio (LR) ≥ 100 were extracted for further analysis, which included 718 viral RBPs and 314 human cell membrane proteins. The RBPs of human-infecting viruses were compiled from three sources: the ViralZone database (Masson et al., 2012), the UniprotKB database in which viral proteins were annotated with GO terms “viral entry into host cell” or “virion attachment the host cell”, and the literatures related to viral RBPs. The viruses belonging to the same viral family were supposed to use the same RBPs. For example, all coronaviruses were supposed to take the spike protein as RBPs.

To evaluate the ability of the RF model in identification of virus receptors, 25 pairs of experimentally validated interactions between viral RBPs and receptors, and the predicted PPIs between these viral RBPs and human cell membrane proteins were extracted from the P-HIPSTer database. For each viral RBP, the predicted RBP-interacting human cell membrane proteins were ranked by either the LR provided in Lasso’s work, or the predicted score provided by the RF model. Then, the ranks of the real receptors were analyzed, and the rank percentage of each real receptor was calculated by dividing the rank by the number of RBP-interacting human cell membrane proteins.

When ranking the RBP-interacting proteins by the RF model, the performance of the RF model may be over-estimated due to the sequence similarity between the RBP-interacting proteins and human proteins in the modeling. To reduce the above effect, for each pair of viral RBP and receptor, the predicted RBP-interacting human cell membrane proteins were clustered with human proteins used in the modeling using CD-HIT at 50% identity level. All the proteins which were clustered with RBP-interacting proteins were excluded in the modeling.

### Data availability

All data used in this study were obtained from public databases as mentioned above.

## Results

### Development of random-forest models for predicting the receptorome of the human-infecting virome

Our previous studies have shown that human virus protein receptors have unique features including high N-glycosylation level, high number of interaction partners in the human PPI network, and high expression level in 32 common human tissues (Zhang et al., 2019). To identify the potential receptors of the human-infecting virome, firstly, a RF model was built to distinguish the human virus receptor proteins from other human membrane proteins based on the above features. The RF model built based on individual protein feature achieved an AUC ranging from 0.51 to 0.61 in five-fold cross-validations (Table 1). The combination of all three features greatly improved the RF model with both the AUC and the prediction accuracy equaling to 0.70 (Table 1).

**Table 1.** The predictive performances of RF models using different sets of features. N-gly, N-glycosylation; PPI, node degree in human PPI network; Expression, expressions in 32 human tissues. Acc, accuracy; Sen, Sensitivity; Spe, Specificity.

| Model with different sets of features | Acc | Sen | Spe | AUC |
| --- | --- | --- | --- | --- |
| N-gly | 0.59 | 0.58 | 0.59 | 0.59 |
| PPI | 0.62 | 0.60 | 0.62 | 0.61 |
| Expression | 0.50 | 0.51 | 0.50 | 0.51 |
| N-gly+PPI+Expression | 0.72 | 0.68 | 0.72 | 0.70 |
| Top 10 k-mers | 0.70 | 0.71 | 0.70 | 0.71 |
| N-gly+PPI+Expression + Top 10 k-mers | 0.76 | 0.73 | 0.76 | 0.75 |

For comparison, we also developed RF models to distinguish the human virus receptors from other human membrane proteins based on protein sequences. The frequencies of k-mers with length of one and two amino acids were used as features in the modeling. The RF model built based on the combination of 20 amino-acid frequencies achieved an AUC of 0.70. When using the frequencies of k-mers with two amino acids as features, only the top N (N = 1 to 400) most important features were used in the modeling to reduce over-fitting. Figure S1 showed that the AUC of the RF models increased rapidly as the increase of N from 1 to 10. Then, it increased slowly or even decreased when more than ten features were used. Therefore, only top ten features were used in the modeling to balance the performance and complexity of the model. The RF model based on top ten features achieved an AUC of 0.71.

To further improve the model for predicting the receptorome of the human-infecting virome, the protein features and the top ten most important k-mers of two amino acids were incorporated in the modeling. The RF model achieved an AUC of 0.75. The prediction accuracy, sensitivity and specificity of the model were 0.76, 0.73 and 0.76, respectively (Table 1). The model combining both the protein features and k-mers was used for further analysis.

### Prediction of the receptorome for the human-infecting virome

Based on the RF model, the receptorome was predicted from human cell membrane proteins. A score ranging from 0 to 1 was assigned to each human cell membrane protein. The proteins with high scores are more likely to be virus receptors. A total of 1380 proteins with scores greater than 0.5 were considered to constitute the receptorome of the human-infecting virome. Table 2 listed top 20 human cell membrane proteins and the relevant scores (for all human cell membrane proteins, please see Table S1).

**Table 2.** Top 20 human cell membrane proteins and their scores assigned by the RF model.

| Gene Name | Protein name | RF score | Gene Name | Protein name | RF score |
| --- | --- | --- | --- | --- | --- |
| LAMP1 | Lysosome-associated membrane glycoprotein 1 | 0.968 | LAMP2 | Lysosome-associated membrane glycoprotein 2 | 0.913 |
| ITGAV | Integrin alpha-V | 0.961 | TYRO3 | Tyrosine-protein kinase receptor TYRO3 | 0.911 |
| PVR | Poliovirus receptor | 0.955 | HSPA8 | Heat shock cognate 71 kDa protein | 0.907 |
| FGFR3 | Fibroblast growth factor receptor 3 | 0.953 | GPNMB | Transmembrane glycoprotein NMB | 0.899 |
| NECTIN1 | Nectin-1 | 0.946 | NCAM1 | Neural cell adhesion molecule 1 | 0.895 |
| TNFRSF14 | Tumor necrosis factor receptor superfamily member 14 | 0.928 | CD86 | T-lymphocyte activation antigen CD86 | 0.893 |
| DPP4 | Dipeptidyl peptidase 4 | 0.928 | EGFR | Epidermal growth factor receptor | 0.893 |
| CD55 | Complement decay-accelerating factor | 0.922 | LDLR | Low-density lipoprotein receptor | 0.892 |
| ITGB1 | Integrin beta-1 | 0.918 | CD44 | CD44 antigen | 0.891 |
| CR2 | Complement receptor type 2 | 0.915 | FGFR2 | Fibroblast growth factor receptor 2 | 0.889 |

### Prediction of virus-receptor interactions for human-infecting viruses

Then, the prediction of virus-receptor interactions was investigated. In the previous study, Lasso et al. predicted 282,528 pairs of PPIs between human and 1,001 human-infecting viruses (Lasso et al., 2019). Based on the study, 9,395 pairs of PPIs between 718 viral RBPs from 693 human-infecting viruses, and 314 human cell membrane proteins were extracted for further analysis (see Table S2). A viral RBP was predicted to interact with 1 to 65 human cell membrane proteins, with a median of 10. For each viral RBP, the RBP-interacting cell membrane proteins were ranked by the score provided by the RF model to select the most likely receptor (Table S2).

To validate the accuracy of the ranking by the RF model, 25 pairs of experimentally validated interactions between viral RBPs and receptors were extracted. For each pair of viral RBP and its receptor, the rank of the real receptor among the predicted RBP-interacting proteins was obtained, and then the related rank percentage was calculated (Materials and Methods). Ten real receptors were ranked in top one by the RF model (Table 3). Besides, nearly 60% (14/25) of real receptors were ranked in top three. On average, the real receptors had a rank percentage of 0.20 among all the RBP-interacting human cell membrane proteins, suggesting that the real receptors would be ranked in the top 20% of all candidates by the RF model.

**Table 3.**
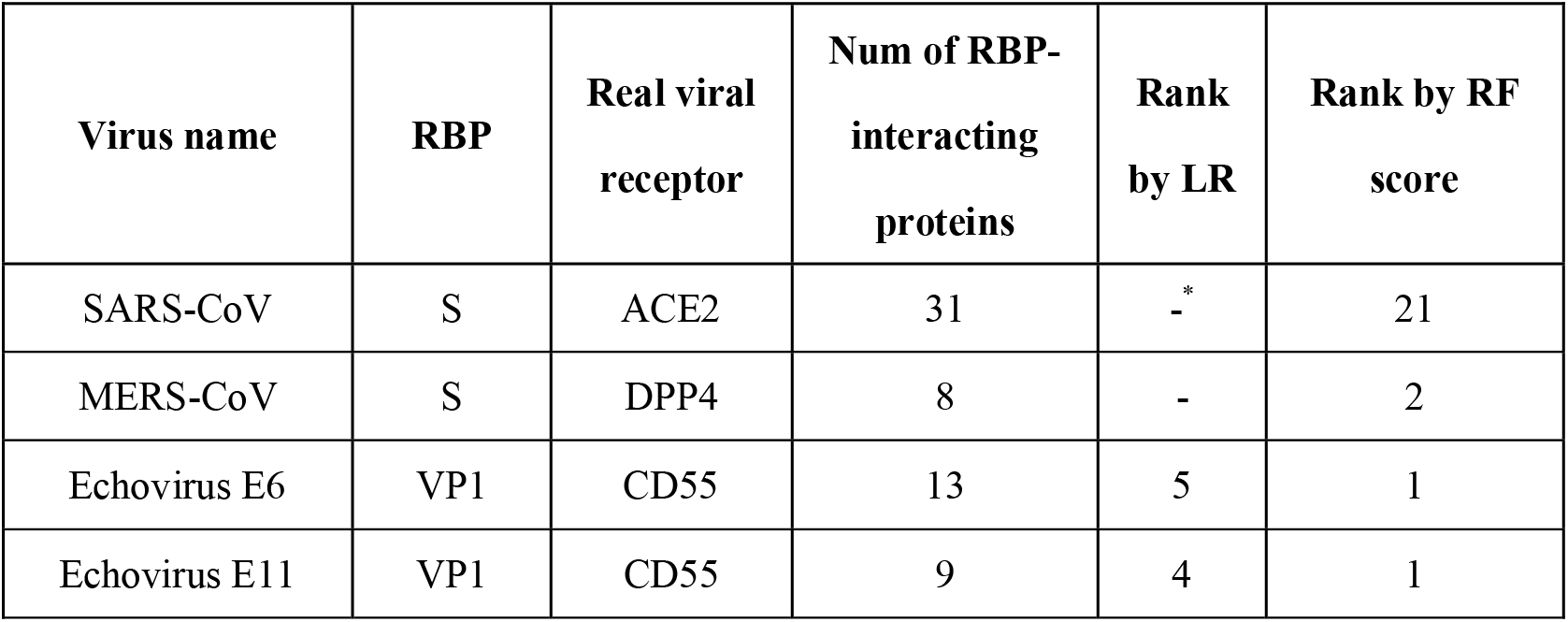
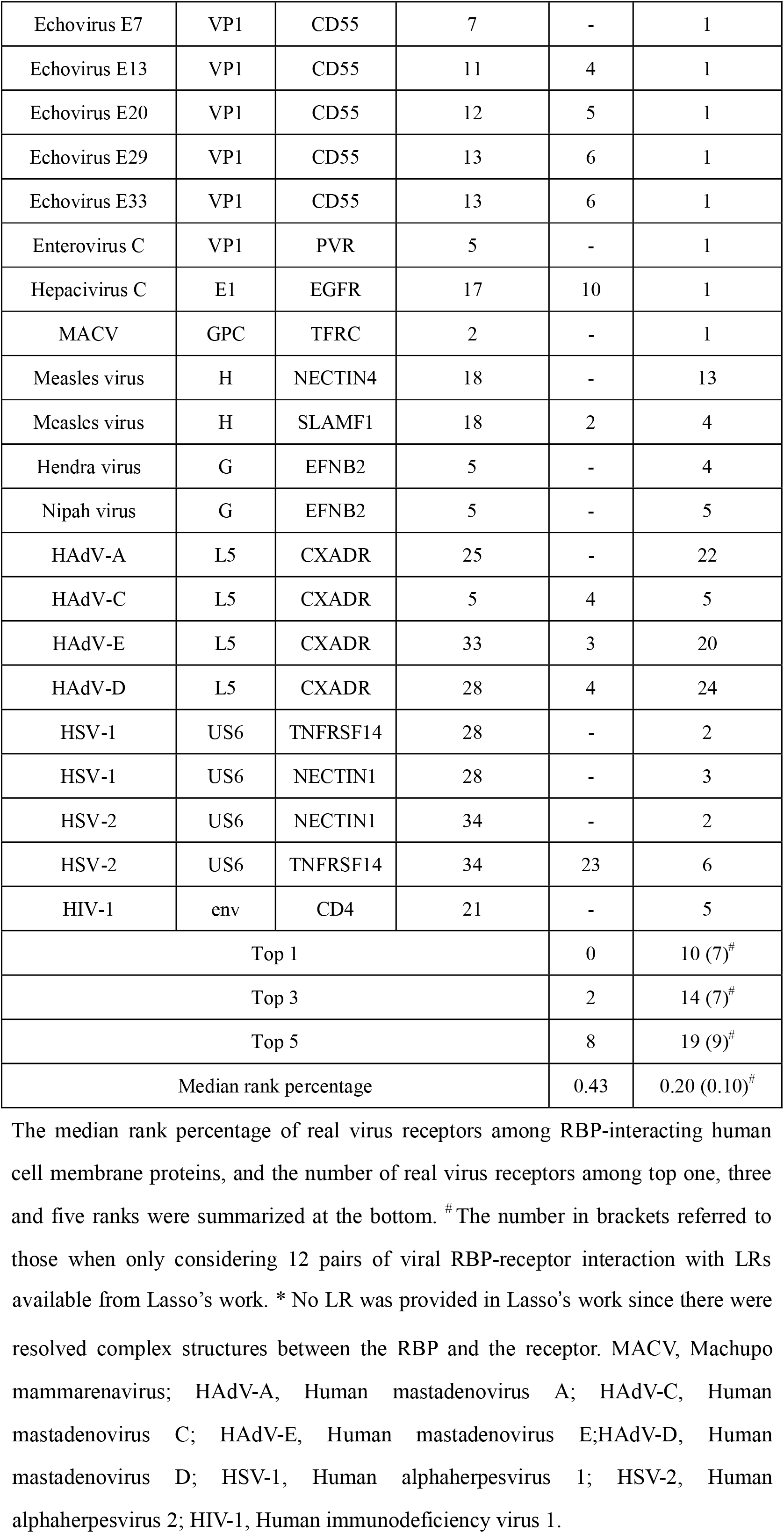
The ranks of real virus receptors among the RBP-interacting human cell membrane proteins by LR and RF score.

| Virus name | RBP | Real viral receptor | Num of RBP-interacting proteins | Rank by LR | Rank by RF score |
| --- | --- | --- | --- | --- | --- |
| SARS-CoV | S | ACE2 | 31 | -* | 21 |
| MERS-CoV | S | DPP4 | 8 | - | 2 |
| Echovirus E6 | VP1 | CD55 | 13 | 5 | 1 |
| Echovirus E11 | VP1 | CD55 | 9 | 4 | 1 |
| Echovirus E7 | VP1 | CD55 | 7 | - | 1 |
| Echovirus E13 | VP1 | CD55 | 11 | 4 | 1 |
| Echovirus E20 | VP1 | CD55 | 12 | 5 | 1 |
| Echovirus E29 | VP1 | CD55 | 13 | 6 | 1 |
| Echovirus E33 | VP1 | CD55 | 13 | 6 | 1 |
| Enterovirus C | VP1 | PVR | 5 | - | 1 |
| Hepacivirus C | E1 | EGFR | 17 | 10 | 1 |
| MACV | GPC | TFRC | 2 | - | 1 |
| Measles virus | H | NECTIN4 | 18 | - | 13 |
| Measles virus | H | SLAMF1 | 18 | 2 | 4 |
| Hendra virus | G | EFNB2 | 5 | - | 4 |
| Nipah virus | G | EFNB2 | 5 | - | 5 |
| HAdV-A | L5 | CXADR | 25 | - | 22 |
| HAdV-C | L5 | CXADR | 5 | 4 | 5 |
| HAdV-E | L5 | CXADR | 33 | 3 | 20 |
| HAdV-D | L5 | CXADR | 28 | 4 | 24 |
| HSV-1 | US6 | TNFRSF14 | 28 | - | 2 |
| HSV-1 | US6 | NECTIN1 | 28 | - | 3 |
| HSV-2 | US6 | NECTIN1 | 34 | - | 2 |
| HSV-2 | US6 | TNFRSF14 | 34 | 23 | 6 |
| HIV-1 | env | CD4 | 21 | - | 5 |
| Top 1 |  |  |  | 0 | 10 (7)<sup>#</sup> |
| Top 3 |  |  |  | 2 | 14 (7)<sup>#</sup> |
| Top 5 |  |  |  | 8 | 19 (9)<sup>#</sup> |
| Median rank percentage |  |  |  | 0.43 | 0.20 (0.10)<sup>#</sup> |

The median rank percentage of real virus receptors among RBP-interacting human cell membrane proteins, and the number of real virus receptors among top one, three and five ranks were summarized at the bottom. <sup>#</sup>The number in brackets referred to those when only considering 12 pairs of viral RBP-receptor interaction with LR<sub>s</sub>
available from Lasso's work. \* No LR was provided in Lasso's work since there were resolved complex structures between the RBP and the receptor. MACV, Machupo mammarynavirus; HAdV-A, Human mastadenovirus A; HAdV-C, Human mastadenovirus C; HAdV-E, Human mastadenovirus E; HAdV-D, Human mastadenovirus D; HSV-1, Human alphaherpesvirus 1; HSV-2, Human alphaherpesvirus 2; HIV-1, Human immunodeficiency virus 1.

The LR provided in Lasso’s work can also be used to rank the RBP-interacting proteins. 12 of 25 pairs of experimentally validated viral RBP-receptor interactions had LRs available from Lasso’s work. For comparison, the viral RBP-interacting human cell membrane proteins were ranked by LR. No real receptor was ranked in top one, and only two real receptors were ranked in top three when ranking RBP-interacting human cell membrane proteins by using LR. On average, the median rank percentage of real receptors was 0.43 when ranking was conducted by the LR, while that was 0.10 by the RF model (Table 3).

## Discussion

The identification of receptors for human-infecting viruses is critical for understanding the interactions between viruses and human. Our previous studies have shown that human virus receptor proteins have some unique features compared to other cell membrane proteins, including high N-glycosylation level, high number of interaction partners and high expression level. This study further built a RF model for identifying human virus receptors from human cell membrane proteins with an accepted accuracy. Based on the RF model, the receptorome for the human-infecting virome was predicted, which included a total of 1380 human cell membrane proteins. The results could facilitate the identification of human virus receptors.

In the previous study, Lasso et al. developed a computational model for predicting PPIs between human-infecting viruses and human (Lasso et al., 2019). A variable number of human cell membrane proteins were predicted to interact with viral RBPs. To further select the potential receptors for viruses, both the LR and the RF model were used to rank the viral RBP-interacting human cell membrane proteins. The RF model was found to rank the real receptors better than the LR in a small validation dataset, suggesting that the performance of the RF model may be superior to that of the LR in selecting the real receptors from the predicted RBP-interacting human cell membrane proteins. The combination of the RF model and the RBP-interacting human cell membrane proteins predicted in Lasso’s work enabled the prediction of receptors for 693 human-infecting viruses (Table S2). Nevertheless, more efforts are needed to validate these candidate receptors in future studies.

There are some limitations to this study. Firstly, the number of human virus receptor proteins was much smaller than that of human membrane proteins in the modeling, which may hinder accurate modeling. Thus, the under-sampling method was used to deal with the imbalance problem. Secondly, the performance of the RF model was modest in discriminating human virus receptor proteins from human membrane proteins. More efforts are still needed to improve the model. Thirdly, although the RF model can be used to predict the receptorome of human-infecting virome, it is not feasible to use the model to identify the receptors for a specific human-infecting virus. The combination of the RF model with the model of PPI predictions such as Lasso’s work can help identify virus-receptor interactions.

In conclusion, this study for the first time built a computational model for predicting the receptorome of the human-infecting virome. The results can facilitate the identification of human virus receptors in either computational or experimental studies.

## Acknowledgements

This work was supported by the National Key Plan for Scientific Research and Development of China (2016YFD0500300), Hunan Provincial Natural Science Foundation of China (2018JJ3039), and the Chinese Academy of Medical Sciences (2016-I2M-1-005).

The authors have declared that no competing interests exist.

## Ethical Statement

No ethical approval was required because no human or animal samples were collected in this study.

