## Supplementary material for "Prediction of the receptorome for the human-infecting virome": Figure S1

**Figure S1**. The AUC of the random-forest model base on the top N (N = 1 to 400) most important features (frequencies of k-mer with two amino acids).

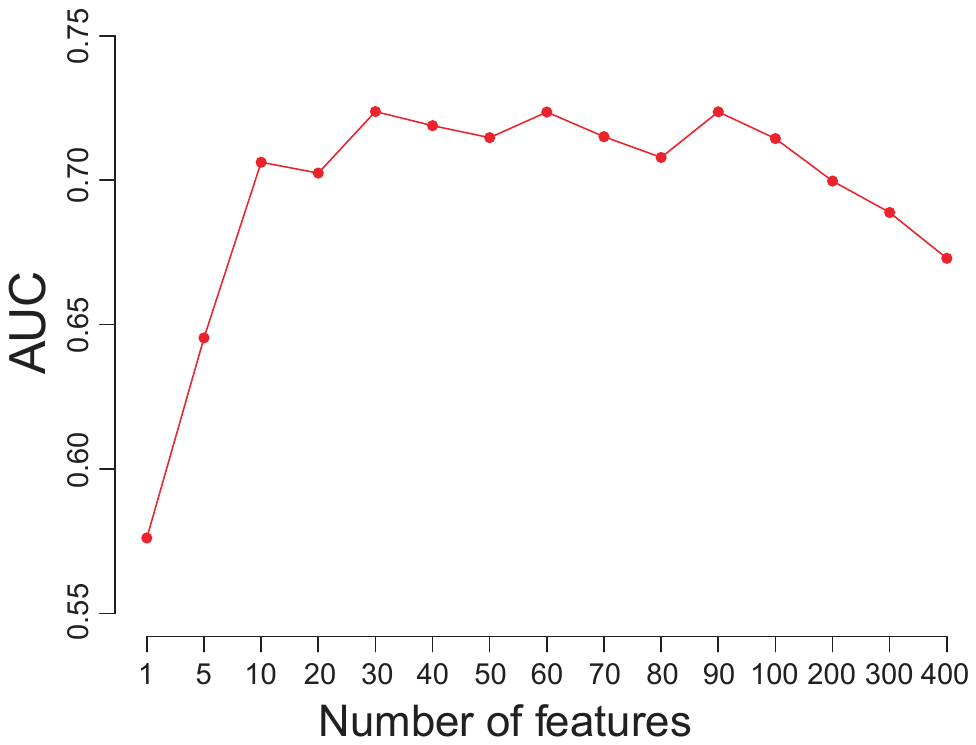
